# Mentorship as a strategy to improve research ability of students and young researchers in Africa: an exploratory study and initial findings on the CORE Africa Research Mentorship Scheme

**DOI:** 10.1101/2021.01.29.428492

**Authors:** Lem Ngongalah, Ngwa Niba Rawlings, James Musisi, Kimonia Awanchiri, Emmanuella Akwah, Etienne Ngeh, Andrew Ssemwanga

## Abstract

Mentorship provides an opportunity to support skill development, improve research ability, promote interest in research and offer career advice. The need for research mentorship in Africa is well-recognised. However, there is scarce literature on the development of such programmes and their potential impacts on students and young researchers in Africa (SYRIA). This study documents the development of the CORE Africa Research Mentorship Scheme (CARMS), and reports the outcomes and challenges experienced over a two-year period, from 2018 to 2020.

41 mentors and mentees from Cameroon, Uganda and Nigeria participated in the programme. Mentors were based in Africa and the UK, while mentees were all based in Africa. Mentees gained knowledge and skills in various research areas including proposal writing, research methods, data analysis, report writing and research publication. Nine mentees gained their first research publication through the CARMS and three successfully completed their first grant proposal. Three mentees were supported by their mentor to develop PhD research proposals and five others gained skills that helped them secure paid research jobs. Eleven mentees have ‘graduated’ from the programme, seven of which are currently enrolled as trainee mentors. Mentees appreciated the opportunity to improve on their research skills and felt that the programme gave them a ‘safe’ environment to freely express their concerns.

Main challenges encountered were: difficulty finding suitable mentors, communication barriers, poorly defined mentorship goals, dealing with mentee’s lack of knowledge/experience of ‘basic’ research concepts and funding limitations. This programme had several positive impacts on the knowledge and skills of mentees and demonstrates the importance of mentorship in research capacity strengthening. Sustaining such programmes requires investments in training and development, to ensure that mentees are continuously and adequately supported.

## 1. Introduction

Mentorship is instrumental for research capacity strengthening, as it provides an opportunity to support skill development, promote interest in research, offer career advice and build professional networks (1). The research mentorship process involves an experienced researcher (the mentor) taking a special interest in guiding another individual – usually a student or another researcher (the mentee) in the development and appraisal of their research ideas, abilities, project activities and professional profiles (2,3). Research mentors can support mentees in a number of ways, including supporting the development of research topics, helping to transform ideas into research projects, planning and managing project activities, seeking research funding and writing and disseminating research reports (3). Mentoring also provides a safe platform where mentees can make and learn from their mistakes, and mentor-mentee relationships foster personal, professional and career advancement.

The need for research mentorship has been identified in various studies across Africa, especially by university students and early career researchers (3–6). The Collaboration for Research Excellence in Africa (CORE Africa) has also identified this need through surveys, interviews and other engagements with students and researchers in Africa (7–10). While much emphasis has been laid on the importance of mentoring to research capacity development (8–11), there is scant literature on the development of such programmes in Africa, and very few studies have explored the potential impacts of mentorship on the research abilities and outcomes of students and young researchers in Africa (SYRIA) (1, 11, 12). Young researchers in the context of this programme refers to researchers who are in their early years of being research-active, typically less than 3 years.

This study describes the structure of the CORE Africa Research Mentorship Scheme (CARMS), detailing how it works, the outcomes on SYRIA between 2018 and 2020, and the challenges faced throughout the development, implementation and delivery of the programme in Cameroon, Uganda and Nigeria.

## 2. Methods

This is an exploratory case study based on CORE Africa’s experience of developing a research mentorship programme for SYRIA.

### 2.1. Development of the CARMS

A CORE Africa forum was held in 2017 to discuss the role of mentorship in strengthening the research abilities of SYRIA. This forum was informed by background findings on research challenges in Africa, research capacity strengthening approaches and experiences from other parts of the world (7, 8). The CARMS was developed in 2017, partly in fulfilment of two of CORE Africa’s objectives, which are to provide opportunities that support the development of research skills for African researchers, and to promote collaboration amongst researchers in Africa.

### 2.2. programme goals and target population

The overall aim of the CARMS programme is to connect SYRIA who require support with conducting research to experienced researchers who can provide guidance and support them through their research activities. Mentors are research-active volunteers who are typically professionals in their fields and have at least three years of demonstrable research experience. Mentees are typically university students, recent graduates or professionals in various fields who are new to research or have limited research experience.

### 2.3. Mentor and mentee requirements

In order to join the pool of mentors or mentees, candidates are required to sign up for the programme through the CORE Africa website (13). Candidates fill a questionnaire providing details about their research needs (for mentees) or expertise (for mentors). Mentees are required to select up to three areas where they need mentorship, while mentors select the areas they are interested in mentoring on. Mentees are normally required to either be currently working on a research project or planning to start working on one. Background checks are conducted to confirm the identity and suitability of mentors and mentees (e.g. academic background, past research activities and roles).

### 2.4. Matching and pairing

The CARMS committee matches potential mentors and mentees based on the mentee’s research needs and mentor’s research interests and expertise. The mentor/mentee selection process is iterative, with feedback from both mentees and potential mentors to ensure that there are no conflicts of interest or conflicts in commitment. Potential candidates are contacted by email and invited for an informal interview to provide more details about what they do, their research experience and what they need help on (for mentees) or what areas they can mentor on (for mentors). Candidates with complete information are then invited to read and agree to the program’s terms and conditions, after which they are added to the mentor/mentee database from where they are paired with a suitable mentor or mentee.

### 2.5. Mentor-mentee engagements and commitments

Mentors and mentees sign an agreement form on enrolment to formalize their mentor-mentee relationship and clarify their mentorship expectations. These expectations include their mentorship goals (based on mentee’s needs and mentor’s expertise), mentorship duration, communication frequency, communication preferences and confidentiality agreements. Mentees are required to provide their mentor-mentee agreement form to the CARMS committee for subsequent reviewing and progress checks. Both parties commit to meeting regularly to work together in realising their mentorship goals.

### 2.6. Mentorship activities

The CARMS programme was designed to be flexible, such that mentors are able use a ‘what works best’ approach to support the development of their mentees. Mentors provide support using various methods including verbal advice through one-to-one conversations or phone calls, online and in-person lectures and presentations, reviewing documents and commenting on drafts, communicating by email and sharing material for supplementary learning. Mentors may also support mentees by linking them with other researchers or professionals within their networks who they believe can be helpful in developing mentees’ skills and professional abilities.

### 2.7. Role of the management team and CARMS committee

The role of the management team is to advertise the programme, elicit participation, identify and invite suitable candidates, process new applications and review programme feedback. The CARMS committee is responsible for matching mentors and mentees, monitoring mentorship activities and assessing programme outcomes.

### 2.8. Programme Evaluation

The CARMS is evaluated based on specific programme metrics (see table 1) developed by the CARMS Committee and reviewed on a quarterly basis. These metrics are evaluated through monitoring and evaluation records and mentorship check-point surveys.

**Table 1:** CARMS programme evaluation metrics.

|  |
| --- |
| 1. Number of mentors and mentees participating in the CARMS<br>2. Number of mentees successfully matched to mentors<br>3. Number of mentors successfully matched to mentees<br>4. Mentor-mentee engagement levels<br>5. Knowledge and skills gained by mentees<br>6. Mentee satisfaction rate<br>7. Number of mentees who complete the mentorship scheme<br>8. Number of mentors and mentees retained in the scheme<br>9. Number of institutions across Africa reached through the mentorship scheme |

#### 2.8.1. Mentorship checkpoints

Check-point times vary depending on the duration of each mentor-mentee relationship, typically after the first month and again every three months until completion.

##### 2.8.1.1. Mentee checkpoint surveys

Mentee checkpoints assess for frequency of contact with mentors, knowledge gained by mentees, skills developed by mentees, level of satisfaction with support received from mentors, number of mentorship goals achieved and estimated progression levels (in %). The survey also collects specific comments from mentees about their mentor, their mentor-mentee relationship and any challenges experienced.

##### 2.8.1.2. Mentor checkpoint surveys

Mentor checkpoints assess for frequency of contact with mentees, level of satisfaction with mentee’s engagement levels, estimated progression levels (in %), specific comments about their mentee or mentor-mentee relationship and any challenges experienced. Checkpoint questions were slightly modified in 2019 to reflect programme updates and improvements.

## 3. Results

### 3.1. Programme Participants

A total of 41 mentors and mentees participated in the CARMS between 2018 and 2020; 14 mentors (9 male, 5 female) and 27 mentees (17 male, 10 female). Mentors and mentees were from three African countries – Cameroon, Uganda and Nigeria. 62% of mentors were based in African countries, while 38% were based in the UK. All mentees were based in Africa. Seven mentees joined the programme in the first year (2018) and 18 joined between 2019 and 2020. Two mentees who joined the programme in 2020 dropped out shortly after due to other commitments that prevented them from following up with their mentorship activities. These two mentees are excluded from the analyses in this report.

### 3.2. Mentorship requests

From 2018 to 2020, mentees requested mentorship on a total of 11 research areas. A majority of mentorship requests were on literature searching using scholarly databases (84%), defining a suitable research topic (60%), scientific writing (60%) and research methods (52%) (table 2). Compared to 2018, there was an increase in the number of mentees requesting mentorship in 10 out of the 11 research areas in 2019/2020 (all areas except referencing).

**Table 2:**
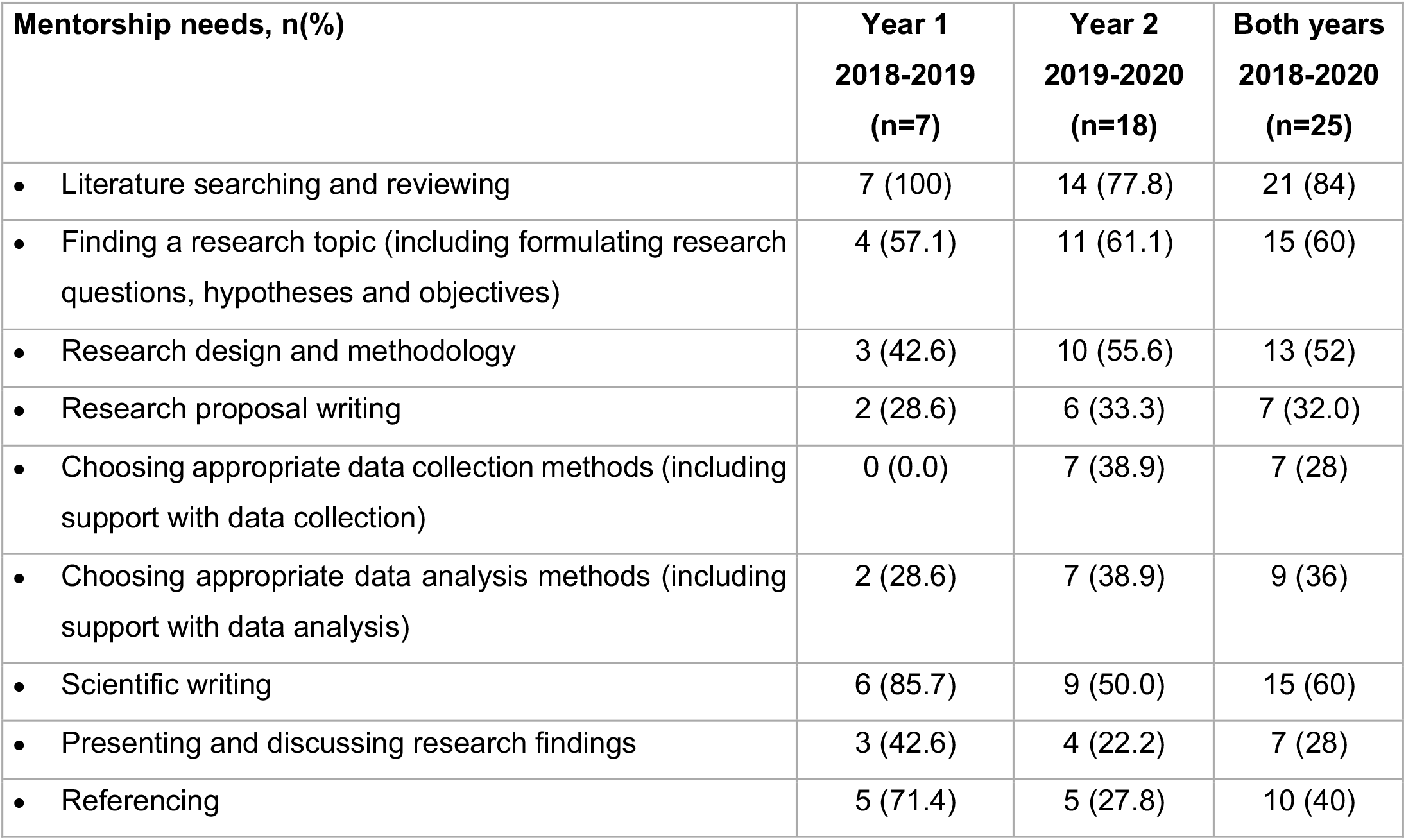
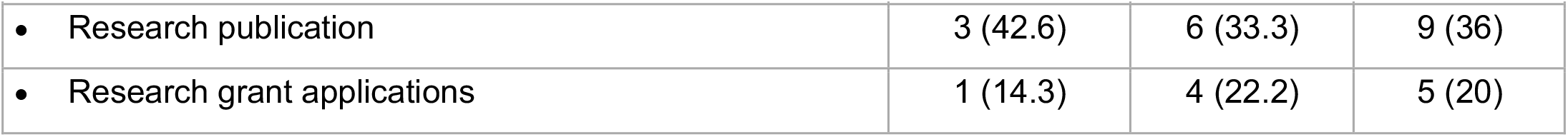
CARMS mentorship requests.

### 3.3. Mentor-mentee engagement levels

On average, a majority of mentees had 2-5 meetings with their mentors within the first month and 5-10 meetings within six months (table 3). About a third of mentees had met with their mentors more than 10 times within a year.

**Table 3:**
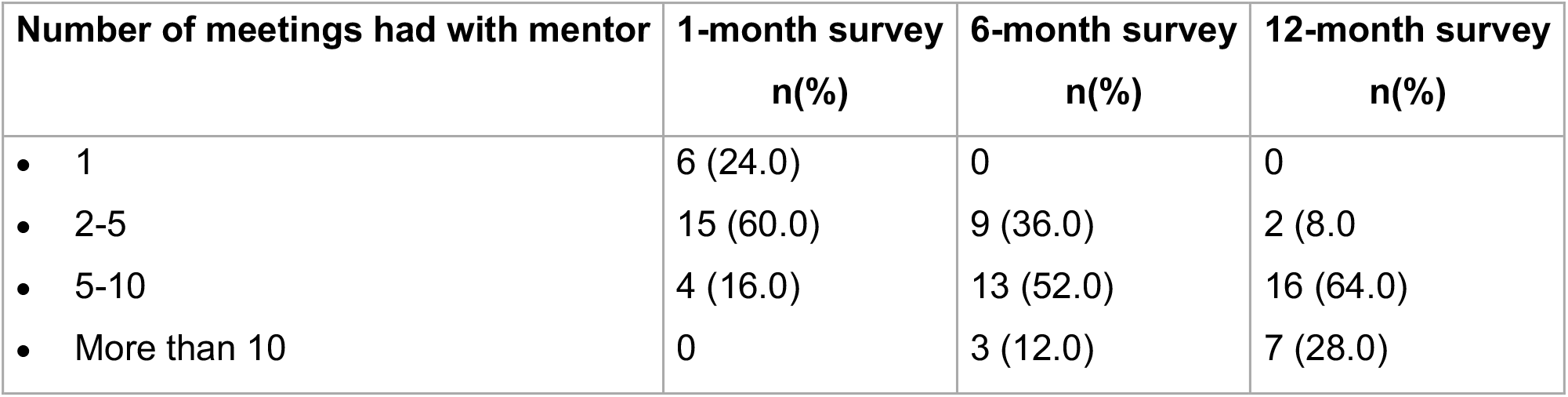
Mentor-mentee engagement levels (average from 2018-2020)

| **Number of meetings had with mentor** | **1-month survey**<br>**n(%)** | **6-month survey**<br>**n(%)** | **12-month survey**<br>**n(%)** |
| --- | --- | --- | --- |
| • 1 | 6 (24.0) | 0 | 0 |
| • 2-5 | 15 (60.0) | 9 (36.0) | 2 (8.0) |
| • 5-10 | 4 (16.0) | 13 (52.0) | 16 (64.0) |
| • More than 10 | 0 | 3 (12.0) | 7 (28.0) |

**Table 4:** Mentee satisfaction and progress (average between 2018-2020)

|  | **1-month**<br>**survey**<br>**n(%)** | **6-month**<br>**survey**<br>**n(%)** | **12-month**<br>**survey**<br>**n(%)** |
| --- | --- | --- | --- |
| **How would you rate the level of support you have received from your mentor?** |  |  |  |
| • Satisfactory | 21 (84) | 23 (92) | 23 (92) |
| • Good | 4 (16.0) | 2 (8.0) | 2 (8.0) |
| • Could be better | 0 | 0 | 0 |
| • Poor | 0 | 0 | 0 |
| **Do you feel you are making progress with your mentorship goals** |  |  |  |
| • Yes | 25 (100) | 25 (100) | 25 (100) |
| • Not really | 0 | 0 | 0 |
| • No | 0 | 0 | 0 |
| **How far would you say you have gone towards achieving your mentorship goals?** |  |  |  |
| • Less than 10% | 6 (24.0) | 2 (8.0) | 0 |
| • 10-30% | 17 (68.0) | 8 (32) | 1 (4.0) |
| • 30-50% | 2 (8.0) | 11 (44) | 4 (16.0) |
| • Over 70% | 0 (0.0) | 2 (8.0) | 9 (36.0) |
| • Completed | 0 (0.0) | 3 (12.0) | 11 (44.0) |

### 3.3. Mentorship outcomes

#### 3.3.1. Knowledge and skills gained by mentees

Figure 1 below summarises the learning outcomes of mentees from 2018-2020. The figure shows the proportion of mentees who requested mentorship in each area (in orange) and the proportion of mentees who reported having gained knowledge or skills in the different mentorship areas (in blue). The number of mentees who reported having gained knowledge or skills in each mentorship area was either equivalent to or higher than the number of mentees who requested mentorship in that area. However, mentees who reported having learnt or gained skills on research design and methodology were slightly less compared to the number of mentorship requests received in those areas. Most mentees also gained knowledge and skills in other research areas besides those for which they requested mentorship – for example, while 84% of mentees signed up for mentorship on literature searching and reviewing, 88% reported having learnt how to do this from their mentor. Mentees also reported having gained knowledge and skills in other aspects that were not included as standard mentorship requests, such as critiquing a research paper and scientific communication.

**Figure 1:**
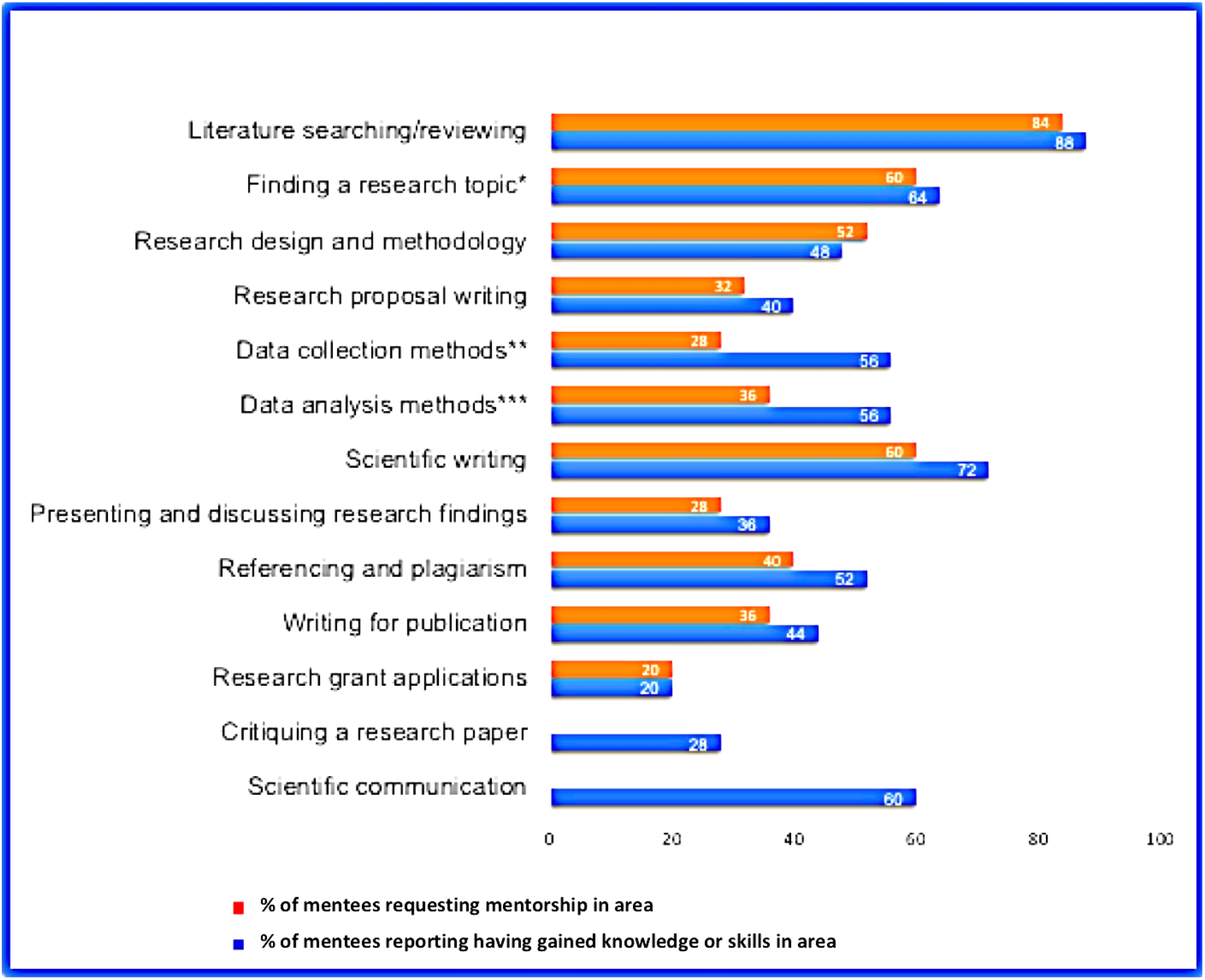
Knowledge and skills gained from the CARMS. **\*Footnote:** *Requests for mentorship on finding a research topic including support on formulating research questions, hypotheses and objectives **Requests for mentorship on choosing appropriate data collection methods including practical support e.g. designing questionnaires, developing interview questions, etc ***Requests for mentorship on analysing different types of data s including practical support e.g. choosing appropriate statistical tests, using statistical software, etc

#### 3.3.2. Academic and professional development

Nine mentees gained their first research publication through the CARMS programme and three successfully completed their first grant proposal. Three mentees were supported by their mentor to develop PhD research proposals and five others found that the knowledge and skills they gained from their mentor helped them secure paid research jobs in their countries.

#### 3.3.3. Mentee retention, satisfaction and progress

The mentee retention rate for this programme is 92.6% due to the two mentees who dropped out as described in section 3.1. All mentees were satisfied with the levels of support provided by their mentors and indicated that they were making progress towards achieving their mentorship goals. A majority of mentees had achieved 10-30% of their learning targets within one month, 30-50% within 3 months and over 70% within 12 months.

Eleven mentees have ‘graduated’ from their formal mentor-mentee relationships. Mentees were considered to have ‘graduated’ when they had achieved their mentorship goals, as delineated at the point of entry and agreed with their mentor. Of the 11 mentees who graduated, seven have been enrolled in a trainee mentorship programme where they are being prepared to mentor others.

#### 3.3.4. Mentees’ perceptions of their mentors and mentorship experience

A majority of mentees rated both their mentor and mentorship experience as ‘great’ (see table 5). Mentees appreciated the opportunity to improve on their research skills and commented about their mentors being ‘knowledgeable’, ‘reliable’, ‘caring’ and ‘flexible’. Mentees also felt that the programme gave them a ‘safe’ environment to freely express their concerns. Sample mentee comments are shown in Appendix 1.

**Table 5:**
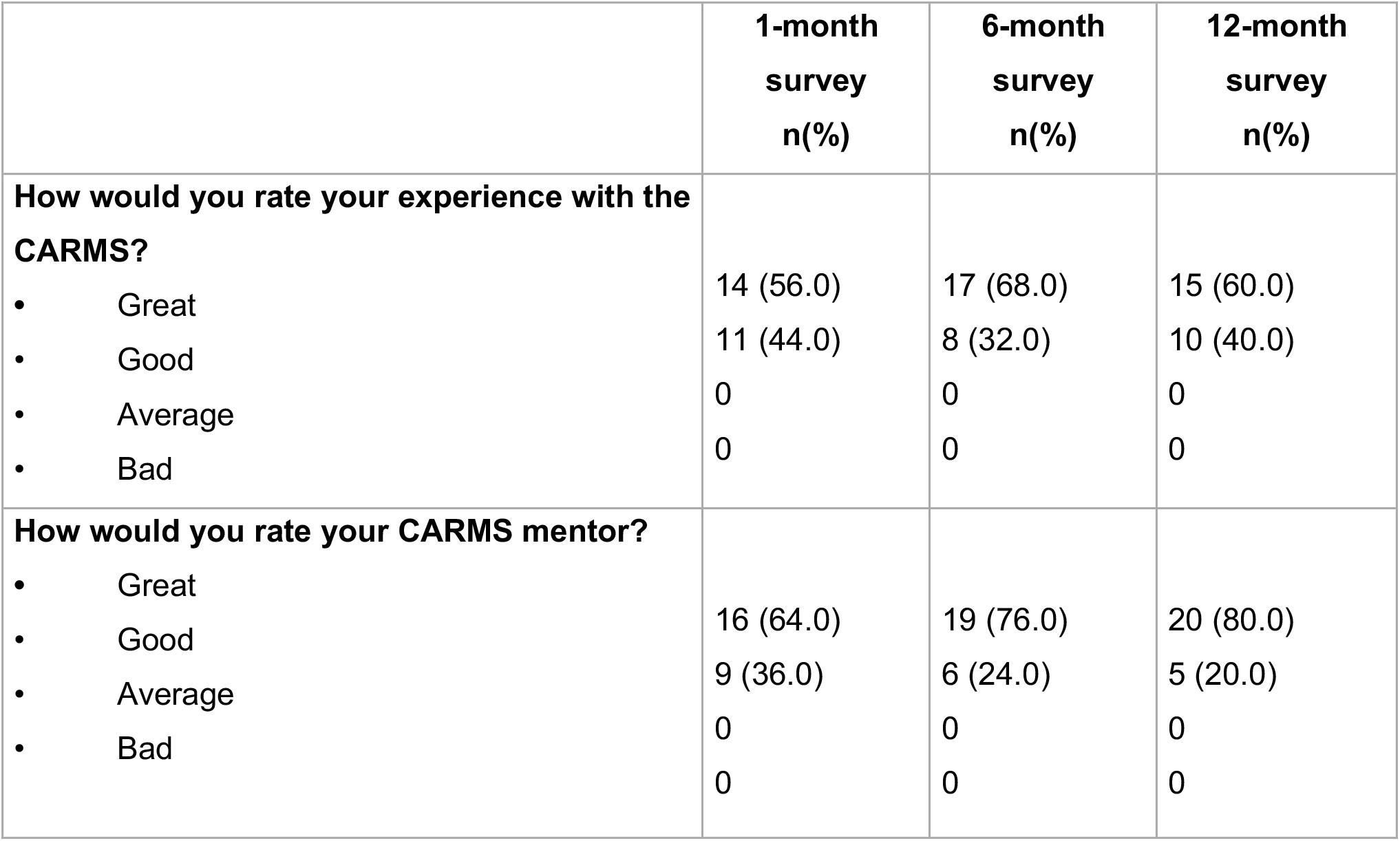
Mentorship experience and perceptions of mentors (average 2018-2020)

|  | **1-month**<br>**survey**<br>**n(%)** | **6-month**<br>**survey**<br>**n(%)** | **12-month**<br>**survey**<br>**n(%)** |
| --- | --- | --- | --- |
| **How would you rate your experience with the CARMS?** |  |  |  |
| • Great | 14 (56.0) | 17 (68.0) | 15 (60.0) |
| • Good | 11 (44.0) | 8 (32.0) | 10 (40.0) |
| • Average | 0 | 0 | 0 |
| • Bad | 0 | 0 | 0 |
| **How would you rate your CARMS mentor?** |  |  |  |
| • Great | 16 (64.0) | 19 (76.0) | 20 (80.0) |
| • Good | 9 (36.0) | 6 (24.0) | 5 (20.0) |
| • Average | 0 | 0 | 0 |
| • Bad | 0 | 0 | 0 |

### 3.4. programme Challenges

Five main challenges were experienced throughout the development, implementation and delivery of the mentorship programme.

#### (i) Finding mentors

One of the most difficult steps during the development of this programme was getting suitable mentors to participate. While several volunteers showed interest in participating and signed up to the program, finding candidates who meet the criteria for mentors and show potential to help mentees succeed was a challenge.

#### (ii) Poorly defined goals

We identified that some mentees did not really know what they wanted to achieve from their mentorship relationship. Although mentees were required to define their mentorship needs and goals when signing up to the program, some mentors found during their first mentor-mentee meetings that their mentees’ goals were ‘unrealistic’ or ‘overambitious’. This was a challenge mostly during the first year of the program.

To mitigate this challenge, mentee goal-setting was included as one of the components to be discussed and agreed upon by both mentors and mentees during their first meeting. This was done to ensure that both parties were satisfied with the targets and expectations of their mentorship.

#### (iii) Communication and internet barriers

Some mentors commented about mentees not responding to emails on time or not providing reasons for failing to respect deadlines. As such, mentors found themselves having to chase after mentees for feedback or updates. While reasons for this may be varied, internet challenges contributed significantly to this problem. Several mentees faced challenges with getting access to their emails due to power cuts or connectivity issues, which sometimes lasted days. These problems were common for mentees who either lived in remote areas or travelled frequently to these areas for work purposes. Communication issues also affected the collection of check-point data from mentees for progress reviews, and this usually required several follow-up calls or messages, resulting in the process taking long to complete.

#### (iv) Dealing with mentees’ lack of knowledge and inexperience

Although all mentors were experienced in their various research fields, concerns were raised about mentors having to teach mentees ‘basic things’ which they expected mentees to have learned during their academic studies – for example, basic research concepts like differences between quantitative and qualitative research, or how to use software like Microsoft Word and Excel. Such challenges were common amongst mentees who were students (mostly Masters or final year undergraduate students working on their academic research projects). This challenge contributed to their mentorship running longer than initially planned. The eligibility criteria for the mentorship programme have since been revised and mentees requiring support on basic research concepts not directly related to their actual projects are supported through another CORE Africa programme (14).

#### (v) Funding limitations

Although all CARMS mentors have a personal desire to support SYRs and are committed to doing so, this requires a significant amount of time and effort on their part. While no mentor has dropped out of the programme since inception, there is a risk of mentors feeling demotivated overtime, as there is no remuneration for their work. This also means that the number of mentees who can benefit from this programme remains limited, as this is subject to the availability of mentors. Sustaining such programmes also requires continuous investments in the training of mentors and development of mentorship platforms. Yet, funding remains a challenge.

## 4. Discussion

This exploratory case study documents the development of a research mentorship programme targeting students and young researchers in Africa, and reports the outcomes and challenges experienced over a two-year period, from 2018 to 2020.

In line with other studies reporting the benefits of research mentorship (1, 3, 4), we have shown that our mentorship programme had a positive influence on the knowledge and skills of mentees. This is especially important in the context of African countries where research challenges abound. Improving the research abilities of students and young researchers can make a significant contribution to increasing research output in Africa, which is currently known to be very low (7, 15). This will also help increase research efforts targeting societal challenges in African countries.

CARMS mentors being based in different African countries and some outside Africa means that part of the programme is delivered virtually. This collaboration across countries facilitates the harmonisation of efforts to support research in Africa. It also provides an opportunity to integrate shared lessons on the research context and challenges in various countries. The positive impacts seen from this programme show that virtual mentorship across African countries can have a positive impact albeit limited by internet and electricity challenges. With the flexible nature of the programme and devotion of both mentors and mentees, mentor-mentee pairs were still able to work together in achieving their goals. Virtual mentorship programs can also prove particularly useful for SYRIA living in conflict-affected countries, where access to physical workshops or research training centres may be limited. In addition, this may be even more important in the era of COVID-19 and current guidelines limiting mobility and physical interaction.

The challenge of having to teach mentees ‘basic things’ during their research mentorship pertained mostly to students, which implies that some students may not have received sufficient support during their academic studies to prepare them for research. This finding aligns with our previous findings on academic research challenges among students in Cameroon and Uganda (9, 10). In light of the benefits reported from this programme, it may be useful for academic institutions to consider establishing such programmes for their students, to enhance research outcomes and output. While some academic institutions may have these in place, considerations need to be made on their effectiveness and potential to meet students’ development needs, especially those not met during their academic studies.

Using a ‘what works best’ approach worked successfully in supporting mentees. However, there were some challenges in unifying programme outcomes especially since mentors used individual mentorship styles and preferences. The CARMS committee has since taken the decision to develop a mentorship handbook to provide an activity guide for both mentors and mentees. This guide is aimed at simplifying the mentorship process for mentors and enhancing mentees’ interaction and learning progress, by defining a range of activities and sample tutorials that can be used during mentorship. This guide takes into consideration what was found to work best in the last two years, as well as input from current mentors. This will serve as a blueprint for the programme, which can be used in addition to individual mentorship preferences.

### Strengths and limitations

This study is the first of its kind to document the development of a research mentorship programme for students and researchers across countries in Africa. It also provides some useful insights on the feasibility and challenges of delivering mentorship virtually in a resource-limited context. The fact that several mentees demonstrated a high level of commitment to supporting other mentees after completing their mentorship shows potential for growth and sustainability. While limited by the small number of participants who have taken part in the programme, our study identifies a number of key findings which require further research, especially considering the scarcity of evidence on this subject.

### Conclusion

The impact of mentorship in research capacity strengthening is well-recognised and the importance of research mentors needs to be acknowledged. Despite the challenges associated with developing and implementing such programmes especially in resource-limited contexts, we have shown that research mentorship can significantly improve the personal and professional development of SYRIA. Ensuring the continuity of such programmes is essential for creating a positive research culture and supportive research environment in Africa. We also note that such programmes require continuous training, development and investments to harness their potential in addressing research challenges and strengthening research capacity in Africa.

## Acknowledgement

We would like to acknowledge the relentless efforts of all mentors whose commitment and dedication made this project possible.

## Appendix 1 Sample comments from mentees about their CARMS mentor and experience

i. “The mentors are well informed about the research process” (Mentee, Nigeria)
ii. “My mentor is very friendly and understanding and this has kept me doing my work without doing it under fear or pressure” (Mentee, Uganda)
iii. “My mentor is very fast in response to email communication. As much as we haven’t had many voice calls, we have exchanged several emails. Recently he reviewed my application for a fellowship. He provided very constructive input and even went ahead to put suggested edits in my concept paper” (Mentee, Uganda)
iv. “I have gained a wider understanding of the research process” (Mentee, Nigeria)
v. “The mentor always reaches out if he notices that I have been silent” (Mentee, Uganda)
vi. “It is amazing how my mentor creates a safe space to freely express my views and interact in a friendly manner” (Mentee, Uganda)
vii. “My mentor is very knowledgeable about how to conduct research” (Mentee, Cameroon)
viii. “I love the flexibility” (Mentee, Cameroon)
ix. “My mentor is very committed. Sometimes when I have very poor connection and we cannot have an online meeting, she’ll spend time talking with me through voice notes on Whatsapp. I’m grateful” (Mentee, Uganda)
x. I don’t usually get enough follow-up from my project supervisors and sometimes it’s not as easy to share my challenges because you have to show independence and there are things they expect you to already know. But with my mentor, I can be honest about what I’m struggling with and it’s nice to see that some of the things I’m talking about, he went through as well. I’ve learnt so much from his experience.

